# Intrinsic Protein Disorder, Conditional Folding and AlphaFold2

**DOI:** 10.1101/2022.03.03.482768

**Authors:** Damiano Piovesan, Alexander Miguel Monzon, Silvio C.E. Tosatto

## Abstract

Intrinsically disordered regions (IDRs) defying the traditional protein structure-function paradigm have been difficult to analyze. AlphaFold’s recent breakthrough in predicting protein structures accurately offers a fresh perspective on IDR prediction as assessed on the CAID dataset. Surprisingly, AlphaFold is highly competitive for predicting both IDRs and conditionally folded regions, demonstrating the plasticity of the disorder to structure continuum.

## Main

The prediction of protein tertiary structure from sequence has been considered the Holy Grail of structural biology since at least the 1960’s, with generations of researchers claiming progress. Since 1994, the biennial Critical Assessment of techniques for protein Structure Prediction (CASP) experiment has tried to measure the state-of-the-art and progress in the field. In CASP14^1^ AlphaFold has at last demonstrated a breakthrough thanks to its clever use of machine learning and multiple sequence alignments ^2,3^. This is leading to a paradigm shift for structural biology due to the sudden availability of orders of magnitude more protein structures ^4,5^. AlphaFoldDB expands the impact further by allowing interested researchers to browse predictions for proteins in several major model organisms^6,7^. This wealth of information is being used to map out less studied parts of the proteome^8,9^. It has highlighted the presence of a considerable fraction of the human proteome with low AlphaFold accuracy scores that may reflect intrinsically disordered regions (IDRs) in proteins ^7,10^. IDRs lacking a fixed tertiary structure are well-known in structural biology^11^ and have been associated to a variety of biological functions^12^.

We have recently described the first round of the Critical Assessment of Protein Intrinsic Disorder (CAID) experiment^13^. CAID aims to establish the state-of-the-art for IDRs in a similar way to CASP, providing two separate challenges. Prediction of IDRs and prediction of those IDR sub-regions responsible for, mostly transient, binding to other molecules. Here, we tested how AlphaFold performs in comparison to other state-of-the-art methods for both categories. From our results, we will draw some interesting conclusions regarding the relationship between IDRs, binding and predicted structures.

Since AlphaFold is a predictor for protein tertiary structure, we first need to establish how it can be used to predict IDRs. The authors of the method already noted how regions for low predicted accuracy (pLDDT score) are correlated to IDRs^6^. In addition to this definition, AlphaFold-pLDDT in the following, we explored another simple definition. Visual inspection shows how AlphaFold tends to predict regions without stable structure as “ribbons” surrounding the folded core (see **Figure 1**). Intuitively, these residues share a high solvent accessible surface, which can be used as a proxy to describe the “ribbon” structure. A simple sliding window of relative accessible surface area (AlphaFold-RSA) has been implemented to benchmark IDR prediction. The size of the sliding window has been optimized based on its performance on the CAID dataset and should be considered an optimistic upper bound for its performance.

**Figure 1.**
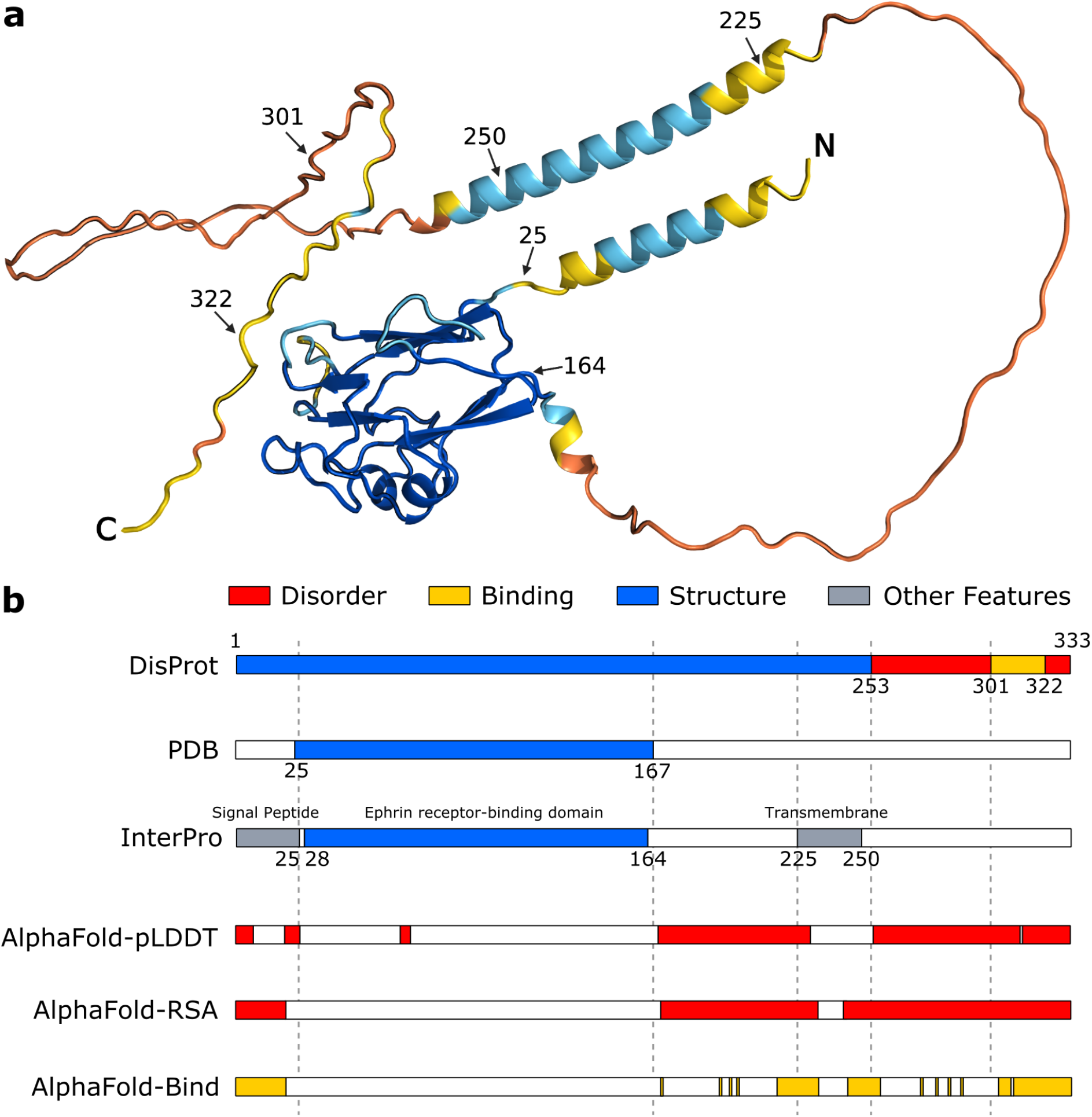
Intrinsically disordered regions and conditional folding derived from AlphaFold prediction. Panel **a,** the structure of the human Ephrin-B2 protein predicted by AlphaFold and colored by pLDDT score (<50 orange, <70 yellow, <90 light blue, >90 dark blue). Residue labels indicate annotated region boundaries. Panel **b**, database annotations (DisProt DP01588, PDB, InterPro P52799) and predicted regions (AlphaFold-pLDDT, AlphaFold-RSA, AlphaFold-Bind). PDB annotation is generated by combining observed residues in different PDB experiments. Per-residue predictions are provided in **Supplementary Figure 1**.

Visual inspection also shows that some “ribbon” regions retain some local secondary structure and have a relatively high pLDDT score. The combination of relative solvent accessibility and pLDDT score can likewise be used to identify regions with a tendency to be simultaneously accessible and structured, which may indicate disordered binding regions or conditional folding (AlphaFold-Bind). For a formal description of the three benchmarked AlphaFold variants, please see the Methods section. **Figure 1** shows an example of the definitions for IDRs and binding regions alongside the predicted AlphaFold tertiary structure. CAID benchmarking is based on DisProt annotations which include IDRs (DisProt) and disordered binding regions (DisProt-binding) datasets, where the negatives are non-annotated residues. To account for incompleteness of IDRs annotation, CAID defines a third dataset, the DisProt-Protein Data Bank (DisProt-PDB), where the negatives are restricted to PDB^14^ Observed residues, so that experimentally ‘uncertain’ residues are excluded.

As expected, AlphaFold-pLDDT performs well on the CAID PDB-DisProt dataset ^6^ (see **Figure 2A, Supp. Tables 1, 2**). The performance is increased using AlphaFold-RSA, which has the highest accuracy of all tested IDR prediction methods. As previously noted, this definition suggests intrinsic disorder can be considered the opposite of globular structure ^13^. When using the DisProt definition, the situation changes somewhat (see **Figure 2B**, **Supp. Tables 3, 4**), with AlphaFold-RSA being among the top five methods. This result can suggest a difference between lack of structure and intrinsic disorder. Indeed when compared to other methods, both AlphaFold-RSA and AlphaFold-pLDDT show a lower precision at low recall (**Figure 2C**) and a lower TPR at low FPR (**Figure 2D**), that is when focusing on the left part of the precision-recall and ROC curves. This indicates that the score does not follow disorder confidence, i.e. high score is not predictive of *bona fide* disorder. When used to identify fully disordered proteins AlphaFold-pLDDT and AlphaFold-RSA behave differently, the first under-predicts and the second over-predicts (**Supp. Table 7**). The former always predicts some residual structure, even if transient.

**Figure 2.**
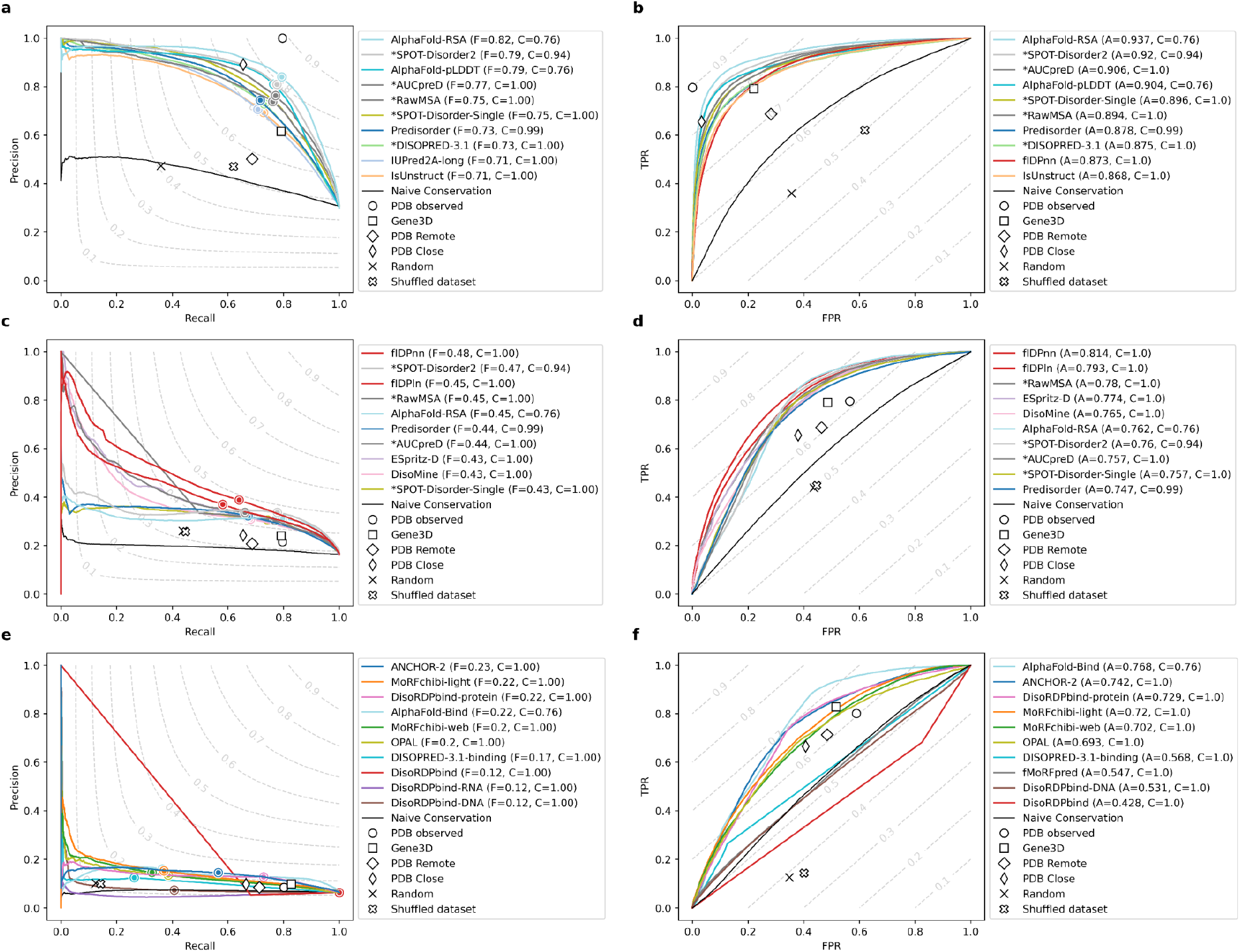
Results for AlphaFold on the three main CAID categories. The results for the DisProt-PDB (n= 646 proteins, panels **A,B**), DisProt (n= 646 proteins, panels **C,D**) and DisProt-binding (n= 646 proteins, panels **E,F**) references are shown. Performance of predictors expressed as maximum F1-Score across all thresholds (F_max_) (panels **A,C,E**) and AUC (panels **B,D,F**) for the top ten best ranking methods (colored) and baselines (white symbols) are shown. The legend on the right of each panel shows the name of the method alongside its F_max_ or AUC score (F and A, respectively) and coverage (C). Notice how the latter is usually 1.0 for most predictors, but only 0.76 for AlphaFold as predictions for some targets are not available.

Conceptually, AlphaFold is not designed to predict binding regions, being limited to single protein chains and intra/protein contacts^3^. When the interaction partner is known and the input properly adapted, AlphaFold has been shown to predict the structure of protein complexes with good accuracy^9^. This is possible because inter/chain contacts between interaction surfaces are well encoded in multiple sequence alignments (and in the model) similarly to intra/chain contacts.

Being able to use AlphaFold to predict disordered binding regions *a priori*, i.e. without knowing the input partner, is therefore not expected. Disorder binding is inherently different from rigid surface-surface interactions, being transient, often multivalent, and versatile, adapting to a number of diverse partners^12^. Indeed, both AlphaFold-pLDDT and AlphaFold-RSA do not perform well on this task (see **Figure 2C, Supp. Tables 5, 6**). Strikingly, the combination of both scores (AlphaFold-Bind) performs very well and reaches state-of-the-art performance on par with ANCHOR2^15^ on the DisProt-Binding dataset.

The ability of AlphaFold to predict IDR binding regions is not entirely surprising. It has been known for a while that binding often occurs in parts of the sequence which have a tendency for disorder close to the decision threshold. This is at the basis of methods such as ANCHOR^16^. Intuitively, the definition we use for AlphaFold falls in the same mold. High solvent accessibility implies the lack of overall structure, while a higher pLDDT score implies some residual local structure. From a biophysical perspective, disorder and secondary structure are both encoded in the protein sequence^17^. As AlphaFold leverages large sequence alignments to generate its models ^3^, it is implicitly encoding sequence conservation in its predictions and this is reflected in the pLDDT score. Hence, it is logical to be able to identify regions undergoing conditional folding.

On the other hand, AlphaFold currently does not provide a thorough description of the structural ensemble for IDRs^10^. The relative movements of residues cannot be captured with a single static structure. AlphaFold-pLDDT fails in identifying fully disordered proteins as some residual structure is always predicted, even if transient (**Supp. Table 7**). Being able to recognize conditional folding events by combining pLDDT and solvent accessibility can however help distinguish in a more coarse grained manner the relative rigidity of the polypeptide chain, separating spacer regions from those involved in transient binding.

Finally, the execution time of AlphaFold is two orders of magnitude slower than methods with similar performance^13^, indicating AlphaFold currently is not the best choice for fast searches at the genomic scale.

We have shown how AlphaFold can be used both to predict IDRs as well as the binding regions inside them. It should be cautioned however that this is not a true blind test and may overestimate its performance. We look forward to being able to assess AlphaFold2 and similar methods fully as part of the next round of CAID.

## Methods

The assessment was carried out exactly as detailed in CAID^13^. The AlphaFoldDB predictions were downloaded via their server on 20 July 2021. Only 489 proteins out of the 645 evaluated in CAID are provided in AlphaFoldDB. The code is implemented as a Python script and freely available from https://github.com/BioComputingUP/AlphaFold-disorder. The three AlphaFold variants are defined as follows.

AlphaFold-pLDDT uses the inverse of the pLDDT score (i.e. 1 - pLDDT) as output. The optimal classification threshold (0.312, representing pLDDT < 68.8%) was selected by maximizing the F1-Score performance on the CAID DisProt dataset.

AlphaFold-RSA calculates the relative solvent accessibility over a local window centered on the residue to predict. The DSSP solvent accessibility^18^ is normalized for each residue by the maximum accessibility of a fully extended Gly-X-Gly peptide^19^ as provided by the BioPython PDB module^20^. The optimal local window size (25 residues, i.e. +/-12), was chosen with a grid search (range: 1 to 50 residues) resulting in a plateau between 20 and 30 residues. Mirroring is used in the local window for positions close to the sequence termini. The optimal classification threshold for AlphaFold-RSA (0.581) was again selected by maximizing the F1-Score performance on the CAID DisProt dataset.

AlphaFold-Bind attempts to recognize binding regions within IDRs through a combination of the previous two scores using the following formula:

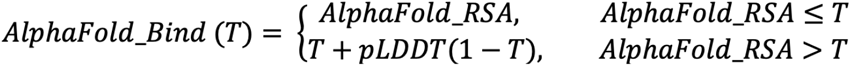

with T set to the AlphaFold-RSA classification threshold (0.581). This ensures values between 0 and 1 by scaling the score for pLDDT accordingly. The optimal classification threshold for AlphaFold-Bind (0.773) was again selected by maximizing the performance on the CAID DisProt-Binding dataset. Notice that this is a proof of principle implementation only. Performance can likely be increased using simple measures such as smoothing over a local window.

## Data availability

The data used including AlphaFold predictions is available from the URL https://idpcentral.org/caid/data/1_alphafold/.

All data used in the analysis are also available in the Code Ocean capsule available at URL https://doi.org/10.24433/CO.6770815.v1.

## Code availability

The CAID assessment can be fully reproduced downloading the code and following the instructions in the CAID GitHub repository at URL https://github.com/BioComputingUP/CAID.

The code is also available and reproducible in the Code Ocean capsule available at URL https://doi.org/10.24433/CO.6770815.v1.

The code to generate AlphaFold disorder predictions is implemented as a Python script and available at URL https://github.com/BioComputingUP/AlphaFold-disorder.

## Acknowledgements

The authors are very grateful to the CAID predictors and DisProt curators for providing the data on which the analysis was made, as well as the AlphaFoldDB developers for making the database openly available. Members of the BioComputing UP lab are acknowledged for insightful discussions. This project has received funding from the European Union’s Horizon 2020 research and innovation programme under the Marie Skłodowska-Curie grant agreement No 778247, the Italian Ministry of University and Research PRIN 2017 grant 2017483NH8 and ELIXIR, the European infrastructure for biological data.

## Author contributions

DP and AMM collected the predictions, produced the data, carried out the assessment and wrote the initial manuscript. SCET designed the experiment, guided the overall project and edited the manuscript. All authors contributed to the discussions and writing of the manuscript.

## Ethics Declaration

The authors declare no competing interests.

