## Supplementary material for "Intrinsic Protein Disorder, Conditional Folding and AlphaFold2": Supp.

*Damiano Piovesan, Alexander Miguel Monzon, Silvio C.E. Tosatto**
*Dept. of Biomedical Sciences, University of Padova, Italy.*

#### Supplementary Information

##### Table of Contents

Fully Disordered Proteins 1
*Supplementary Table 1. Confusion matrix and metrics for the prediction of fully disordered proteins in the DisProt dataset 2*
*Supplementary Table 2. AlphaFold-RSA performance of predicted fully disordered proteins 4*

DisProt-PDB dataset 4
*Supplementary Table 3. Per-residue classification performance in the DisProt-PDB dataset 5*
*Supplementary Table 4. Per-protein classification performance in the DisProt-PDB dataset 6*

DisProt-dataset 6
*Supplementary Table 5. Per-residue classification performance in the DisProt dataset 7*
*Supplementary Table 6. Per-protein classification performance in the DisProt dataset 8*

DisProt-Binding dataset 8
*Supplementary Table 7. Per-residue classification performance in the DisProt-Binding dataset 9*
*Supplementary Table 8. Per-protein classification performance in the DisProt-Binding dataset 9*

Figures 9

#### DisProt-PDB dataset

| | **BAC** | **F1-s** | **FPR** | **MCC** | **PPV** | **TPR** | **TNR** | **COV** |
|---|---|---|---|---|---|---|---|---|
| **PDB observed** | **0.898** | **0.886** | **0.000** | **0.854** | **1.000** | **0.796** | **1.000** | **646** |
| *AlphaFold-RSA* | *0.862* | *0.816* | *0.058* | *0.742* | *0.851* | *0.783* | *0.942* | *489* |
| *SPOT-Disorder2 | 0.848 | 0.792 | 0.078 | 0.705 | 0.811 | 0.773 | 0.922 | 610 |
| *AlphaFold-pLDDT* | *0.842* | *0.786* | *0.070* | *0.701* | *0.821* | *0.753* | *0.930* | *489* |
| *AUCpreD | 0.831 | 0.767 | 0.108 | 0.660 | 0.763 | 0.771 | 0.892 | 644 |
| **PDB Close** | **0.811** | **0.755** | **0.033** | **0.689** | **0.891** | **0.655** | **0.967** | **604** |
| *SPOT-Disorder-Single | 0.822 | 0.753 | 0.118 | 0.640 | 0.743 | 0.763 | 0.882 | 646 |
| *RawMSA | 0.819 | 0.749 | 0.119 | 0.635 | 0.740 | 0.758 | 0.881 | 646 |
| AUCpreD-np | 0.804 | 0.731 | 0.115 | 0.612 | 0.738 | 0.724 | 0.885 | 646 |
| Predisorder | 0.802 | 0.730 | 0.108 | 0.612 | 0.748 | 0.712 | 0.892 | 642 |
| *DISOPRED-3.1 | 0.802 | 0.729 | 0.108 | 0.612 | 0.746 | 0.712 | 0.892 | 646 |
| fIDPnn | 0.792 | 0.710 | 0.150 | 0.574 | 0.686 | 0.735 | 0.850 | 645 |
| IsUnstruct | 0.790 | 0.710 | 0.133 | 0.579 | 0.706 | 0.714 | 0.867 | 646 |

| | | | | | | | | |
|---|---|---|---|---|---|---|---|---|
| VSL2B | 0.789 | 0.707 | 0.139 | 0.574 | 0.696 | 0.718 | 0.861 | 644 |
| IUPred2A-long | 0.785 | 0.706 | 0.110 | 0.582 | 0.733 | 0.680 | 0.890 | 646 |
| MobiDB-lite | 0.783 | 0.704 | 0.106 | 0.583 | 0.739 | 0.673 | 0.894 | 645 |
| IUPred2A-short | 0.787 | 0.704 | 0.142 | 0.569 | 0.692 | 0.717 | 0.858 | 646 |
| ESpritz-X | 0.783 | 0.699 | 0.138 | 0.563 | 0.694 | 0.704 | 0.862 | 645 |
| DisoMine | 0.782 | 0.695 | 0.167 | 0.550 | 0.661 | 0.732 | 0.833 | 646 |
| ESpritz-D | 0.784 | 0.694 | 0.199 | 0.542 | 0.633 | 0.767 | 0.801 | 645 |
| **Gene3D** | **0.785** | **0.692** | **0.220** | **0.539** | **0.615** | **0.791** | **0.780** | **646** |
| fIDPln | 0.777 | 0.687 | 0.183 | 0.536 | 0.642 | 0.738 | 0.817 | 645 |
| ESpritz-N | 0.767 | 0.681 | 0.121 | 0.546 | 0.707 | 0.656 | 0.879 | 645 |
| JRONN | 0.771 | 0.681 | 0.163 | 0.532 | 0.659 | 0.704 | 0.837 | 645 |
| DynaMine | 0.753 | 0.656 | 0.192 | 0.490 | 0.618 | 0.698 | 0.808 | 645 |
| PyHCA | 0.749 | 0.652 | 0.180 | 0.488 | 0.627 | 0.678 | 0.820 | 646 |
| FoldUnfold | 0.736 | 0.636 | 0.193 | 0.462 | 0.608 | 0.666 | 0.807 | 621 |
| DisEMBL-465 | 0.709 | 0.598 | 0.216 | 0.406 | 0.566 | 0.633 | 0.784 | 644 |
| *S2D-2 | 0.706 | 0.596 | 0.292 | 0.385 | 0.517 | 0.704 | 0.708 | 644 |
| **PDB Remote** | **0.703** | **0.579** | **0.282** | **0.376** | **0.501** | **0.687** | **0.718** | **523** |
| DisEMBL-HL | 0.640 | 0.536 | 0.481 | 0.262 | 0.413 | 0.762 | 0.519 | 644 |
| GlobPlot | 0.656 | 0.535 | 0.294 | 0.295 | 0.479 | 0.605 | 0.706 | 645 |
| *DisPredict-2 | 0.620 | 0.514 | 0.469 | 0.222 | 0.403 | 0.708 | 0.531 | 646 |
| **Conservation** | **0.614** | **0.508** | **0.473** | **0.212** | **0.398** | **0.701** | **0.527** | **646** |
| DFLpred | 0.498 | 0.471 | 1.000 | -0.043 | 0.308 | 0.996 | 0.000 | 646 |

**Supplementary Table 1. Per-residue classification performance in the *DisProt-PDB* dataset.** Performance of predictors and baselines for the *DisProt-PDB* dataset. Metrics are averaged over the whole dataset, sorted by F1-Score and predictor thresholds are optimized on F1-Score. Baselines are shown in bold and AlphaFold in italics and underlined. The evaluation metrics are as follows: balanced accuracy (BAC), F1-Score (F1-s), false positive rate (FPR); Matthews correlation coefficient (MCC), positive predictive value (PPV, i.e. precision), true positive rate (TPR, i.e. recall), true negative rate (TNR, i.e. specificity) and coverage (COV), i.e. number of predicted target proteins (out of 646).

| | **BAC** | **F1-s** | **FPR** | **MCC** | **PPV** | **TPR** | **TNR** | **COV** |
|---|---|---|---|---|---|---|---|---|
| **PDB observed** | **0.817** | **0.751** | **0.260** | **0.426** | **0.847** | **0.844** | **0.740** | **646** |
| *AlphaFold-pLDDT* | *0.766* | *0.655* | *0.104* | *0.398* | *0.734* | *0.678* | *0.896* | *489* |
| *AlphaFold-RSA* | *0.769* | *0.659* | *0.142* | *0.380* | *0.745* | *0.690* | *0.858* | *489* |
| *SPOT-Disorder2 | 0.776 | 0.666 | 0.140 | 0.358 | 0.728 | 0.707 | 0.860 | 610 |
| **PDB Close** | **0.716** | **0.624** | **0.291** | **0.341** | **0.780** | **0.712** | **0.709** | **604** |
| *AUCpreD | 0.770 | 0.656 | 0.174 | 0.336 | 0.691 | 0.726 | 0.826 | 644 |
| *DISOPRED-3.1 | 0.719 | 0.597 | 0.132 | 0.310 | 0.702 | 0.621 | 0.868 | 646 |
| Predisorder | 0.727 | 0.607 | 0.170 | 0.299 | 0.669 | 0.658 | 0.830 | 642 |

| | | | | | | | | |
|---|---|---|---|---|---|---|---|---|
| *RawMSA | 0.746 | 0.623 | 0.198 | 0.299 | 0.683 | 0.690 | 0.802 | 646 |
| AUCpreD-np | 0.738 | 0.617 | 0.183 | 0.296 | 0.663 | 0.678 | 0.817 | 646 |
| *SPOT-Disorder-Single | 0.743 | 0.620 | 0.207 | 0.287 | 0.664 | 0.690 | 0.793 | 646 |
| MobiDB-lite | 0.707 | 0.582 | 0.177 | 0.268 | 0.656 | 0.621 | 0.823 | 645 |
| ESpritz-X | 0.722 | 0.594 | 0.220 | 0.268 | 0.634 | 0.677 | 0.780 | 645 |
| IUPred2A-short | 0.724 | 0.598 | 0.211 | 0.268 | 0.641 | 0.671 | 0.789 | 646 |
| IsUnstruct | 0.717 | 0.597 | 0.213 | 0.264 | 0.643 | 0.661 | 0.787 | 646 |
| VSL2B | 0.709 | 0.589 | 0.214 | 0.262 | 0.641 | 0.651 | 0.786 | 644 |
| ESpritz-N | 0.690 | 0.555 | 0.165 | 0.260 | 0.646 | 0.594 | 0.835 | 645 |
| JRONN | 0.705 | 0.577 | 0.213 | 0.254 | 0.632 | 0.646 | 0.787 | 645 |
| **Gene3D** | **0.740** | **0.625** | **0.380** | **0.243** | **0.657** | **0.778** | **0.620** | **646** |
| fIDPnn | 0.729 | 0.607 | 0.311 | 0.243 | 0.630 | 0.731 | 0.689 | 645 |
| IUPred2A-long | 0.685 | 0.548 | 0.184 | 0.241 | 0.664 | 0.585 | 0.816 | 646 |
| DynaMine | 0.698 | 0.565 | 0.233 | 0.236 | 0.603 | 0.647 | 0.767 | 645 |
| fIDPln | 0.718 | 0.593 | 0.361 | 0.219 | 0.623 | 0.740 | 0.639 | 645 |
| PyHCA | 0.682 | 0.543 | 0.258 | 0.215 | 0.606 | 0.632 | 0.742 | 646 |
| DisoMine | 0.696 | 0.563 | 0.333 | 0.205 | 0.618 | 0.686 | 0.667 | 646 |
| DisEMBL-465 | 0.659 | 0.530 | 0.265 | 0.197 | 0.576 | 0.613 | 0.735 | 644 |
| *S2D-2 | 0.666 | 0.526 | 0.317 | 0.191 | 0.561 | 0.654 | 0.683 | 644 |
| **PDB Remote** | **0.664** | **0.521** | **0.349** | **0.185** | **0.595** | **0.651** | **0.651** | **523** |
| FoldUnfold | 0.665 | 0.553 | 0.330 | 0.176 | 0.618 | 0.637 | 0.670 | 621 |
| GlobPlot | 0.622 | 0.486 | 0.305 | 0.152 | 0.549 | 0.576 | 0.695 | 645 |
| ESpritz-D | 0.684 | 0.552 | 0.415 | 0.151 | 0.569 | 0.712 | 0.585 | 645 |
| DisEMBL-HL | 0.663 | 0.533 | 0.533 | 0.131 | 0.513 | 0.771 | 0.467 | 644 |
| **Conservation** | **0.607** | **0.468** | **0.432** | **0.129** | **0.538** | **0.636** | **0.568** | **646** |
| *DisPredict-2 | 0.621 | 0.496 | 0.600 | 0.053 | 0.495 | 0.723 | 0.400 | 646 |
| DFLpred | 0.645 | 0.549 | 0.999 | -0.001 | 0.470 | 0.997 | 0.001 | 646 |

**Supplementary Table 2. Per-protein classification performance in the *DisProt-PDB* dataset**
Performance of predictors and baselines for *DisProt-PDB* dataset. Metrics are averaged over the whole dataset, sorted by F1-Score and predictor thresholds are optimized on F1-Score. Baselines are shown in bold and AlphaFold in italics and underlined. The evaluation metrics are as follows: balanced accuracy (BAC), F1-Score (F1-s), false positive rate (FPR); Matthews correlation coefficient (MCC), positive predictive value (PPV, i.e. precision), true positive rate (TPR, i.e. recall), true negative rate (TNR, i.e. specificity) and coverage (COV), i.e. number of predicted target proteins (out of 646).

#### DisProt-dataset

| | BAC | F1-s | FPR | MCC | PPV | TPR | TNR | COV |
|---|---|---|---|---|---|---|---|---|
| fIDPnn | 0.719 | 0.483 | 0.186 | 0.369 | 0.394 | 0.624 | 0.814 | 645 |
| *SPOT-Disorder2 | 0.722 | 0.470 | 0.325 | 0.345 | 0.338 | 0.769 | 0.675 | 610 |
| fIDPln | 0.691 | 0.452 | 0.179 | 0.329 | 0.378 | 0.561 | 0.821 | 645 |

| | | | | | | | | |
|---|---|---|---|---|---|---|---|---|
| *AlphaFold-RSA* | *0.723* | *0.447* | *0.332* | *0.336* | *0.313* | *0.779* | *0.668* | *489* |
| *RawMSA | 0.703 | 0.446 | 0.248 | 0.323 | 0.338 | 0.654 | 0.752 | 646 |
| *AUCpreD | 0.709 | 0.436 | 0.350 | 0.315 | 0.304 | 0.768 | 0.650 | 644 |
| Predisorder | 0.686 | 0.435 | 0.257 | 0.298 | 0.332 | 0.629 | 0.743 | 642 |
| *SPOT-Disorder-Single | 0.701 | 0.433 | 0.298 | 0.308 | 0.313 | 0.700 | 0.702 | 646 |
| ESpritz-D | 0.682 | 0.431 | 0.215 | 0.301 | 0.343 | 0.578 | 0.785 | 645 |
| *AlphaFold-pLDDT* | *0.701* | *0.426* | *0.341* | *0.303* | *0.298* | *0.744* | *0.659* | *489* |
| AUCpreD-np | 0.688 | 0.426 | 0.270 | 0.295 | 0.317 | 0.647 | 0.730 | 646 |
| DisoMine | 0.691 | 0.425 | 0.288 | 0.295 | 0.311 | 0.669 | 0.712 | 646 |
| IUPred2A-short | 0.683 | 0.420 | 0.269 | 0.287 | 0.314 | 0.634 | 0.731 | 646 |
| MobiDB-lite | 0.678 | 0.420 | 0.236 | 0.288 | 0.326 | 0.592 | 0.764 | 645 |
| IsUnstruct | 0.682 | 0.418 | 0.276 | 0.284 | 0.310 | 0.640 | 0.724 | 646 |
| ESpritz-X | 0.686 | 0.418 | 0.299 | 0.286 | 0.303 | 0.672 | 0.701 | 645 |
| IUPred2A-long | 0.681 | 0.417 | 0.278 | 0.283 | 0.309 | 0.640 | 0.722 | 646 |
| VSL2B | 0.678 | 0.409 | 0.303 | 0.273 | 0.296 | 0.659 | 0.697 | 644 |
| JRONN | 0.661 | 0.401 | 0.242 | 0.260 | 0.311 | 0.563 | 0.758 | 645 |
| ESpritz-N | 0.663 | 0.400 | 0.262 | 0.258 | 0.303 | 0.587 | 0.738 | 645 |
| *DISOPRED-3.1 | 0.671 | 0.394 | 0.370 | 0.255 | 0.272 | 0.712 | 0.630 | 646 |
| PyHCA | 0.655 | 0.386 | 0.309 | 0.239 | 0.280 | 0.619 | 0.691 | 646 |
| DynaMine | 0.652 | 0.385 | 0.288 | 0.237 | 0.285 | 0.592 | 0.712 | 645 |
| **Gene3D** | **0.653** | **0.368** | **0.486** | **0.226** | **0.240** | **0.791** | **0.514** | **646** |
| DisEMBL-465 | 0.633 | 0.367 | 0.255 | 0.214 | 0.283 | 0.520 | 0.745 | 644 |
| FoldUnfold | 0.642 | 0.365 | 0.382 | 0.211 | 0.251 | 0.666 | 0.618 | 621 |
| **PDB Close** | **0.637** | **0.353** | **0.380** | **0.202** | **0.242** | **0.655** | **0.620** | **604** |
| *S2D-2 | 0.623 | 0.347 | 0.423 | 0.181 | 0.234 | 0.668 | 0.577 | 644 |
| **PDB observed** | **0.616** | **0.339** | **0.565** | **0.174** | **0.215** | **0.796** | **0.435** | **646** |
| DisEMBL-HL | 0.605 | 0.333 | 0.312 | 0.162 | 0.244 | 0.522 | 0.688 | 644 |
| *DisPredict-2 | 0.601 | 0.329 | 0.368 | 0.152 | 0.231 | 0.569 | 0.632 | 646 |
| GlobPlot | 0.592 | 0.321 | 0.415 | 0.137 | 0.219 | 0.600 | 0.585 | 645 |
| **PDB Remote** | **0.611** | **0.318** | **0.464** | **0.159** | **0.207** | **0.687** | **0.536** | **523** |
| **Conservation** | **0.547** | **0.294** | **0.682** | **0.075** | **0.181** | **0.776** | **0.318** | **646** |
| DFLpred | 0.498 | 0.279 | 0.998 | -0.032 | 0.162 | 0.994 | 0.002 | 646 |

**Supplementary Table 3. Per-residue classification performance in the *DisProt* dataset**
Performance of predictors and baselines for *DisProt* dataset. Metrics are averaged over the whole dataset, sorted by F1-Score and predictor thresholds are optimized on F1-Score. Baselines are shown in bold and AlphaFold in italics and underlined. The evaluation metrics are as follows: balanced accuracy (BAC), F1-Score (F1-s), false positive rate (FPR); Matthews correlation coefficient (MCC), positive predictive value (PPV, i.e. precision), true positive rate (TPR, i.e. recall), true negative rate (TNR, i.e. specificity) and coverage (COV), i.e. number of predicted target proteins (out of 646).

| | BAC | F1-s | FPR | MCC | PPV | TPR | TNR | COV |
|---|---|---|---|---|---|---|---|---|
| *SPOT-Disorder2 | 0.704 | 0.481 | 0.306 | 0.306 | 0.455 | 0.702 | 0.694 | 610 |
| <u>*AlphaFold-RSA*</u> | <u>*0.689*</u> | <u>*0.455*</u> | <u>*0.317*</u> | <u>*0.303*</u> | <u>*0.444*</u> | <u>*0.685*</u> | <u>*0.683*</u> | <u>*489*</u> |
| <u>*AlphaFold-pLDDT*</u> | <u>*0.691*</u> | <u>*0.435*</u> | <u>*0.283*</u> | <u>*0.303*</u> | <u>*0.423*</u> | <u>*0.667*</u> | <u>*0.717*</u> | <u>*489*</u> |
| *AUCpreD | 0.698 | 0.461 | 0.340 | 0.281 | 0.417 | 0.723 | 0.660 | 644 |
| *RawMSA | 0.677 | 0.428 | 0.237 | 0.281 | 0.455 | 0.586 | 0.763 | 646 |
| *DISOPRED-3.1 | 0.664 | 0.416 | 0.289 | 0.262 | 0.429 | 0.620 | 0.711 | 646 |
| Predisorder | 0.664 | 0.422 | 0.237 | 0.260 | 0.433 | 0.567 | 0.763 | 642 |
| IUPred2A-short | 0.665 | 0.415 | 0.250 | 0.252 | 0.414 | 0.583 | 0.750 | 646 |
| AUCpreD-np | 0.662 | 0.425 | 0.269 | 0.251 | 0.423 | 0.592 | 0.731 | 646 |
| fIDPnn | 0.667 | 0.439 | 0.304 | 0.247 | 0.417 | 0.622 | 0.696 | 645 |
| *SPOT-Disorder-Single | 0.663 | 0.428 | 0.298 | 0.247 | 0.417 | 0.616 | 0.702 | 646 |
| MobiDB-lite | 0.654 | 0.407 | 0.227 | 0.244 | 0.420 | 0.538 | 0.773 | 645 |
| IsUnstruct | 0.656 | 0.417 | 0.268 | 0.243 | 0.414 | 0.581 | 0.732 | 646 |
| ESpritz-X | 0.666 | 0.425 | 0.316 | 0.240 | 0.391 | 0.642 | 0.684 | 645 |
| IUPred2A-long | 0.649 | 0.388 | 0.244 | 0.239 | 0.433 | 0.542 | 0.756 | 646 |
| VSL2B | 0.652 | 0.410 | 0.280 | 0.237 | 0.407 | 0.582 | 0.720 | 644 |
| ESpritz-N | 0.645 | 0.387 | 0.228 | 0.236 | 0.416 | 0.528 | 0.772 | 645 |
| JRONN | 0.638 | 0.384 | 0.208 | 0.234 | 0.431 | 0.495 | 0.792 | 645 |
| fIDPln | 0.645 | 0.406 | 0.290 | 0.219 | 0.421 | 0.566 | 0.710 | 645 |
| DynaMine | 0.638 | 0.386 | 0.252 | 0.219 | 0.397 | 0.537 | 0.748 | 645 |
| DisoMine | 0.637 | 0.408 | 0.364 | 0.207 | 0.401 | 0.620 | 0.636 | 646 |
| PyHCA | 0.636 | 0.383 | 0.300 | 0.201 | 0.382 | 0.570 | 0.700 | 646 |
| **PDB Close** | **0.598** | **0.383** | **0.365** | **0.199** | **0.404** | **0.561** | **0.635** | **604** |
| **Gene3D** | **0.630** | **0.405** | **0.474** | **0.188** | **0.380** | **0.709** | **0.526** | **646** |
| DisEMBL-465 | 0.617 | 0.368 | 0.256 | 0.180 | 0.370 | 0.498 | 0.744 | 644 |
| *S2D-2 | 0.616 | 0.362 | 0.383 | 0.172 | 0.338 | 0.617 | 0.617 | 644 |
| **PDB Remote** | **0.601** | **0.338** | **0.412** | **0.171** | **0.343** | **0.614** | **0.588** | **523** |
| FoldUnfold | 0.620 | 0.386 | 0.382 | 0.169 | 0.383 | 0.607 | 0.618 | 621 |
| **PDB observed** | **0.609** | **0.428** | **0.507** | **0.164** | **0.408** | **0.697** | **0.493** | **646** |
| ESpritz-D | 0.610 | 0.358 | 0.321 | 0.155 | 0.381 | 0.525 | 0.679 | 645 |
| DisEMBL-HL | 0.605 | 0.346 | 0.342 | 0.137 | 0.323 | 0.551 | 0.658 | 644 |
| GlobPlot | 0.592 | 0.338 | 0.381 | 0.136 | 0.325 | 0.572 | 0.619 | 645 |
| **Conservation** | **0.571** | **0.330** | **0.585** | **0.111** | **0.306** | **0.719** | **0.415** | **646** |
| *DisPredict-2 | 0.560 | 0.319 | 0.484 | 0.058 | 0.293 | 0.579 | 0.516 | 646 |
| DFLpred | 0.532 | 0.348 | 0.996 | 0.000 | 0.257 | 0.995 | 0.004 | 646 |

**Supplementary Table 4. Per-protein classification performance in the *DisProt* dataset**
Performance of predictors and baselines for *DisProt* dataset. Metrics are averaged over the whole dataset, sorted by F1-Score and predictor thresholds are optimized on F1-Score. Baselines are shown in bold and AlphaFold in italics and underlined. The evaluation metrics are as follows: balanced accuracy

(BAC), F1-Score (F1-s), false positive rate (FPR); Matthews correlation coefficient (MCC), positive predictive value (PPV, i.e. precision), true positive rate (TPR, i.e. recall), true negative rate (TNR, i.e. specificity) and coverage (COV), i.e. number of predicted target proteins (out of 646).

#### DisProt-Binding dataset

| | BAC | F1-s | FPR | MCC | PPV | TPR | TNR | COV |
|---|---|---|---|---|---|---|---|---|
| ANCHOR-2 | 0.665 | 0.231 | 0.217 | 0.189 | 0.146 | 0.548 | 0.783 | 646 |
| *AlphaFold-Bind* | *0.617* | *0.222* | *0.123* | *0.164* | *0.161* | *0.357* | *0.877* | *489* |
| DisoRDPbind-protein | 0.690 | 0.216 | 0.330 | 0.194 | 0.127 | 0.711 | 0.670 | 646 |
| MoRFchibi-light | 0.609 | 0.215 | 0.122 | 0.154 | 0.157 | 0.340 | 0.878 | 644 |
| MoRFchibi-web | 0.600 | 0.202 | 0.128 | 0.139 | 0.146 | 0.327 | 0.872 | 644 |
| OPAL | 0.610 | 0.200 | 0.165 | 0.139 | 0.135 | 0.385 | 0.835 | 644 |
| **Gene3D** | **0.656** | **0.175** | **0.516** | **0.153** | **0.098** | **0.828** | **0.484** | **646** |
| DISOPRED-3.1-binding | 0.569 | 0.169 | 0.125 | 0.099 | 0.124 | 0.263 | 0.875 | 646 |
| **PDB observed** | **0.606** | **0.152** | **0.589** | **0.106** | **0.084** | **0.801** | **0.411** | **646** |
| fMoRFpred | 0.526 | 0.125 | 0.587 | 0.026 | 0.069 | 0.639 | 0.413 | 646 |
| DisoRDPbind-DNA | 0.528 | 0.124 | 0.352 | 0.028 | 0.073 | 0.407 | 0.648 | 646 |
| DisoRDPbind-RNA | 0.499 | 0.119 | 0.998 | -0.006 | 0.063 | 0.997 | 0.002 | 646 |
| DisoRDPbind | 0.500 | 0.119 | 1.000 | 0.000 | 0.063 | 1.000 | 0.000 | 646 |

**Supplementary Table 5. Per-residue classification performance in the *DisProt-Binding* dataset**
Performance of predictors and baselines for *DisProt-Binding* dataset. Metrics are averaged over the whole dataset, sorted by F1-Score and predictor thresholds are optimized on F1-Score. Baselines are shown in bold and AlphaFold in italics and underlined. The evaluation metrics are as follows: balanced accuracy (BAC), F1-Score (F1-s), false positive rate (FPR); Matthews correlation coefficient (MCC), positive predictive value (PPV, i.e. precision), true positive rate (TPR, i.e. recall), true negative rate (TNR, i.e. specificity) and coverage (COV), i.e. number of predicted target proteins (out of 646).

| | BAC | F1-s | FPR | MCC | PPV | TPR | TNR | COV |
|---|---|---|---|---|---|---|---|---|
| **Gene3D** | **0.529** | **0.143** | **0.523** | **0.053** | **0.125** | **0.261** | **0.477** | **646** |
| DisoRDPbind-protein | 0.663 | 0.131 | 0.334 | 0.059 | 0.142 | 0.215 | 0.666 | 646 |
| DisoRDPbind | 0.194 | 0.131 | 1.000 | 0.000 | 0.100 | 0.358 | 0.000 | 646 |
| DisoRDPbind-RNA | 0.195 | 0.131 | 0.999 | -0.001 | 0.100 | 0.357 | 0.001 | 646 |
| **PDB observed** | **0.490** | **0.128** | **0.569** | **-0.011** | **0.115** | **0.224** | **0.431** | **646** |
| OPAL | 0.604 | 0.121 | 0.360 | 0.029 | 0.122 | 0.166 | 0.640 | 644 |
| fMoRFpred | 0.430 | 0.116 | 0.627 | 0.004 | 0.100 | 0.236 | 0.373 | 646 |
| *AlphaFold-Bind* | *0.728* | *0.114* | *0.201* | *0.053* | *0.139* | *0.141* | *0.799* | *489* |
| ANCHOR-2 | 0.734 | 0.111 | 0.209 | 0.054 | 0.142 | 0.160 | 0.791 | 646 |

| | | | | | | | | |
|---|---|---|---|---|---|---|---|---|
| MoRFchibi-web | 0.668 | 0.109 | 0.262 | 0.038 | 0.131 | 0.137 | 0.738 | 644 |
| MoRFchibi-light | 0.673 | 0.106 | 0.252 | 0.036 | 0.131 | 0.131 | 0.748 | 644 |
| DISOPRED-3.1-binding | 0.725 | 0.095 | 0.172 | 0.036 | 0.130 | 0.105 | 0.828 | 646 |
| DisoRDPbind-DNA | 0.588 | 0.092 | 0.369 | 0.002 | 0.100 | 0.143 | 0.631 | 646 |

**Supplementary Table 6. Per-protein classification performance in the *DisProt-Binding* dataset** Performance of predictors and baselines for *DisProt-Binding* dataset. Metrics are averaged over the whole dataset, sorted by F1-Score and predictor thresholds are optimized on F1-Score. Baselines are shown in bold and AlphaFold in italics and underlined. The evaluation metrics are as follows: balanced accuracy (BAC), F1-Score (F1-s), false positive rate (FPR); Matthews correlation coefficient (MCC), positive predictive value (PPV, i.e. precision), true positive rate (TPR, i.e. recall), true negative rate (TNR, i.e. specificity) and coverage (COV), i.e. number of predicted target proteins (out of 646).

#### Fully Disordered Proteins

| | TN | FP | FN | TP | MCC | F1-s | TNR | TPR | PPV | BAC |
|---|---|---|---|---|---|---|---|---|---|---|
| fIDPnn | 458 | 8 | 7 | 16 | 0.665 | 0.681 | 0.983 | 0.696 | 0.667 | 0.839 |
| SPOT-Disorder-Single | 463 | 3 | 12 | 11 | 0.599 | 0.595 | 0.994 | 0.478 | 0.786 | 0.736 |
| RawMSA | 453 | 13 | 8 | 15 | 0.569 | 0.588 | 0.972 | 0.652 | 0.536 | 0.812 |
| Predisorder | 460 | 6 | 11 | 12 | 0.572 | 0.585 | 0.987 | 0.522 | 0.667 | 0.754 |
| ESpritz-N | 464 | 2 | 14 | 9 | 0.553 | 0.529 | 0.996 | 0.391 | 0.818 | 0.694 |
| SPOT-Disorder2 | 448 | 18 | 9 | 14 | 0.488 | 0.509 | 0.961 | 0.609 | 0.438 | 0.785 |
| IUPred2A-long | 462 | 4 | 14 | 9 | 0.504 | 0.500 | 0.991 | 0.391 | 0.692 | 0.691 |
| AUCpreD | 459 | 7 | 13 | 10 | 0.485 | 0.500 | 0.985 | 0.435 | 0.588 | 0.710 |
| DisoMine | 437 | 29 | 6 | 17 | 0.491 | 0.493 | 0.938 | 0.739 | 0.370 | 0.838 |
| IsUnstruct | 458 | 8 | 13 | 10 | 0.470 | 0.488 | 0.983 | 0.435 | 0.556 | 0.709 |
| VSL2B | 448 | 18 | 11 | 12 | 0.426 | 0.453 | 0.961 | 0.522 | 0.400 | 0.742 |
| IUPred2A-short | 465 | 1 | 16 | 7 | 0.504 | 0.452 | 0.998 | 0.304 | 0.875 | 0.651 |
| MobiDB-lite | 465 | 1 | 16 | 7 | 0.504 | 0.452 | 0.998 | 0.304 | 0.875 | 0.651 |
| fIDPln | 444 | 22 | 10 | 13 | 0.425 | 0.448 | 0.953 | 0.565 | 0.371 | 0.759 |
| *AlphaFold-RSA* | *427* | *39* | *6* | *17* | *0.436* | *0.430* | *0.916* | *0.739* | *0.304* | *0.828* |
| JRONN | 463 | 3 | 16 | 7 | 0.446 | 0.424 | 0.994 | 0.304 | 0.700 | 0.649 |
| DisPredict-2 | 462 | 4 | 16 | 7 | 0.422 | 0.412 | 0.991 | 0.304 | 0.636 | 0.648 |
| ESpritz-X | 464 | 2 | 17 | 6 | 0.428 | 0.387 | 0.996 | 0.261 | 0.750 | 0.628 |
| ESpritz-D | 435 | 31 | 11 | 12 | 0.340 | 0.364 | 0.933 | 0.522 | 0.279 | 0.728 |
| S2D-2 | 443 | 23 | 13 | 10 | 0.325 | 0.357 | 0.951 | 0.435 | 0.303 | 0.693 |
| **Gene3D** | **401** | **65** | **4** | **19** | **0.385** | **0.355** | **0.861** | **0.826** | **0.226** | **0.843** |
| PyHCA | 464 | 2 | 18 | 5 | 0.380 | 0.333 | 0.996 | 0.217 | 0.714 | 0.607 |
| **PDB observed** | **373** | **93** | **7** | **16** | **0.252** | **0.242** | **0.800** | **0.696** | **0.147** | **0.748** |
| FoldUnfold | 358 | 108 | 6 | 17 | 0.246 | 0.230 | 0.768 | 0.739 | 0.136 | 0.754 |

| | | | | | | | | | | |
|---|---|---|---|---|---|---|---|---|---|---|
| **PDB Remote** | **455** | **11** | **19** | **4** | **0.185** | **0.211** | **0.976** | **0.174** | **0.267** | **0.575** |
| DISOPRED-3.1 | 462 | 4 | 20 | 3 | 0.217 | 0.200 | 0.991 | 0.130 | 0.429 | 0.561 |
| DisEMBL-HL | 466 | 0 | 21 | 2 | 0.288 | 0.160 | 1.000 | 0.087 | 1.000 | 0.543 |
| DisEMBL-465 | 466 | 0 | 22 | 1 | 0.204 | 0.083 | 1.000 | 0.043 | 1.000 | 0.522 |
| **Conservation** | **328** | **138** | **20** | **3** | **-0.077** | **0.037** | **0.704** | **0.130** | **0.021** | **0.417** |
| DFLpred | 466 | 0 | 23 | 0 | 0.000 | 0.000 | 1.000 | 0.000 | 0.000 | 0.500 |
| DynaMine | 466 | 0 | 23 | 0 | 0.000 | 0.000 | 1.000 | 0.000 | 0.000 | 0.500 |
| *AlphaFold-pLDDT* | *466* | *0* | *23* | *0* | *0.000* | *0.000* | *1.000* | *0.000* | *0.000* | *0.500* |
| GlobPlot | 466 | 0 | 23 | 0 | 0.000 | 0.000 | 1.000 | 0.000 | 0.000 | 0.500 |
| **PDB Close** | **456** | **10** | **23** | **0** | **-0.032** | **0.000** | **0.979** | **0.000** | **0.000** | **0.489** |

**Supplementary Table 7. Confusion matrix and metrics for the prediction of fully disordered proteins in the DisProt dataset**

Proteins with disorder prediction or disorder annotation covering at least 95% of the sequence are considered fully disordered. The subset of proteins predicted by AlphaFold are considered (n= 489). Predictors are sorted by their F1-score. Baseline names are in bold. The evaluation metrics are as follows: true negatives (TN(, true positives (TP), false negatives (FN), false positives (FP), F1-score (F1-s), true negative rate (TNR, i.e. specificity) true positive rate (TPR, i.e. recall), positive predictive value (PPV, i.e. precision), balanced accuracy for prediction of fully disordered proteins (BAC).

| Pred | DisProt | UniProt | Protein name | Organism | Length | Content |
|---|---|---|---|---|---|---|
| TP | DP01135 | P40019 | Histone H2A.Z-specific chaperone CHZ1 | *S. cerevisiae* | 153 | 100% |
| TP | DP01159 | A6NF83 | Nuclear protein 2 | *Homo sapiens* | 97 | 100% |
| TP | DP01299 | Q96273 | Late embryogenesis abundant protein 18 | *A. thaliana* | 97 | 100% |
| TP | DP01300 | Q9SLJ2 | Dehydrin HIRD11 | *A. thaliana* | 98 | 100% |
| TP | DP01425 | Q15004 | PCNA-associated factor | *Homo sapiens* | 111 | 100% |
| TP | DP01435 | O60829 | P antigen family member 4 | *Homo sapiens* | 102 | 100% |
| TP | DP01471 | P36118 | Pre-mRNA-splicing factor NTR2 | *S. cerevisiae* | 322 | 100% |
| TP | DP01521 | Q5RJL0 | Ermin | *R. norvegicus* | 282 | 100% |
| TP | DP01559 | Q04964 | Ribonucleotide reductase inhibitor protein SML1 | *S. cerevisiae* | 104 | 100% |
| TP | DP01677 | P06454 | Prothymosin alpha | *Homo sapiens* | 111 | 100% |
| TP | DP01776 | Q8GYN5 | RPM1-interacting protein 4 | *A. thaliana* | 211 | 100% |
| TP | DP01858 | Q96270 | Late embryogenesis abundant protein 7 | *A. thaliana* | 169 | 100% |
| TP | DP01876 | P52926 | High mobility group protein HMGI-C | *Homo sapiens* | 109 | 100% |
| TP | DP02005 | P47710 | Alpha-S1-casein | *Homo sapiens* | 185 | 100% |
| TP | DP02066 | Q9FG31 | Late embryogenesis abundant protein 46 | *A. thaliana* | 158 | 100% |
| TP | DP02078 | P02686-5 | Isoform 5 of Myelin basic protein | *Homo sapiens* | 171 | 100% |
| TP | DP02205 | Q2TUW1 | Abscisic stress ripening-like protein | *Glycine max* | 238 | 100% |
| FP | DP01448 | P10923 | Osteopontin | *Mus musculus* | 294 | 94.56% |
| FP | DP01558 | Q9XTY3 | Seven B Two (Mammalian 7BT prohormone convertase chaperone) homolog | *C. elegans* | 211 | 92.42% |

| | | | | | | |
|---|---|---|---|---|---|---|
| FP | DP01557 | P27682 | Neuroendocrine protein 7B2 | *R. norvegicus* | 210 | 88.57% |
| FP | DP01173 | Q15726 | Metastasis-suppressor KiSS-1 | *Homo sapiens* | 138 | 86.23% |
| FP | DP01473 | Q96GU1 | P antigen family member 5 | *Homo sapiens* | 130 | 85.38% |
| FP | DP01502 | P33328 | Synaptobrevin homolog 2 | *S. cerevisiae* | 115 | 79.13% |
| FP | DP01126 | O43561 | Linker for activation of T-cells family member 1 | *Homo sapiens* | 262 | 78.63% |
| FP | DP01191 | O14140 | 26S proteasome complex subunit rpn15 | *S. pombe* | 71 | 76.06% |
| FP | DP01453 | Q8NHZ8 | Anaphase-promoting complex subunit CDC26 | *Homo sapiens* | 85 | 70.59% |
| FP | DP01246 | O13916 | Anaphase-promoting complex subunit hcn1 | *S. pombe* | 80 | 70.00% |
| FP | DP01658 | Q09882 | Pre-mRNA-processing protein 45 | *S. pombe* | 557 | 61.22% |
| FP | DP01943 | Q9NRY2 | SOSS complex subunit C | *Homo sapiens* | 104 | 59.62% |
| FP | DP01392 | P53330 | Regulator of Ty1 transposition protein 102 | *S. cerevisiae* | 157 | 54.78% |
| FP | DP01454 | P60006 | Anaphase-promoting complex subunit 15 | *Homo sapiens* | 121 | 53.72% |
| FP | DP01498 | O70480 | Vesicle-associated membrane protein 4 | *Mus musculus* | 141 | 50.35% |
| FP | DP01747 | P48788 | Troponin I, fast skeletal muscle | *Homo sapiens* | 182 | 47.25% |
| FP | DP01281 | Q9JM54 | Phorbol-12-myristate-13-acetate-induced protein 1 | *Mus musculus* | 103 | 46.60% |
| FP | DP01455 | Q96DE5 | Anaphase-promoting complex subunit 16 | *Homo sapiens* | 110 | 46.36% |
| FP | DP02296 | P60761 | Neurogranin | *Mus musculus* | 78 | 43.59% |
| FP | DP02269 | Q9Y6K9 | NF-kappa-B essential modulator | *Homo sapiens* | 419 | 40.57% |
| FP | DP01172 | O43516 | WAS/WASL-interacting protein family member 1 | *Homo sapiens* | 503 | 39.17% |
| FP | DP01643 | O75506 | Heat shock factor-binding protein 1 | *Homo sapiens* | 76 | 35.53% |
| FP | DP01608 | Q9Y6H6 | Potassium voltage-gated channel subfamily E member 3 | *Homo sapiens* | 103 | 34.95% |
| FP | DP01883 | P02655 | Apolipoprotein C-II | *Homo sapiens* | 101 | 25.74% |
| FP | DP02061 | P70447 | Neurogenin-2 | *Mus musculus* | 263 | 22.05% |
| FP | DP01203 | O75807 | Protein phosphatase 1 regulatory subunit 15A | *Homo sapiens* | 674 | 17.66% |
| FP | DP01794 | Q99ML1 | Bcl-2-binding component 3 | *Mus musculus* | 193 | 17.62% |
| FP | DP01434 | P31431 | Syndecan-4 | *Homo sapiens* | 198 | 14.14% |
| FP | DP02170 | O15234 | Protein CASC3 | *Homo sapiens* | 703 | 12.94% |
| FP | DP01313 | Q9N4U5 | Smu-2 suppressor of mec-8 and unc-52 protein | *C. elegans* | 547 | 11.15% |
| FP | DP02149 | Q24570 | Cell death protein Grim | *D. melanogaster* | 138 | 10.87% |
| FP | DP01999 | Q8K4J6 | Myocardin-related transcription factor A | *Mus musculus* | 964 | 9.96% |
| FP | DP01139 | A0A0G2JXC5 | Myocardin-related transcription factor A | *R. norvegicus* | 1038 | 9.25% |
| FP | DP02122 | Q02199 | Nucleoporin NUP49/NSP49 | *S. cerevisiae* | 472 | 7.20% |
| FP | DP01603 | Q96IZ0 | PRKC apoptosis WT1 regulator protein | *Homo sapiens* | 340 | 6.76% |
| FP | DP01994 | P27321 | Calpastatin | *R. norvegicus* | 713 | 5.89% |
| FP | DP01366 | Q24139 | Protein commissureless 1 | *D. melanogaster* | 370 | 2.97% |
| FP | DP01278 | Q68DK7 | Male-specific lethal 1 homolog | *Homo sapiens* | 614 | 1.79% |

**Supplementary Table 8. AlphaFold-RSA performance of predicted fully disordered proteins.** The columns are (from left to right): AlphaFold-RSA prediction (Pred; TP: true positive, FP: false positive), DisProt and UniProt accession numbers, protein name and organism, length in amino acids, DisProt disorder content (Content).

### Figures

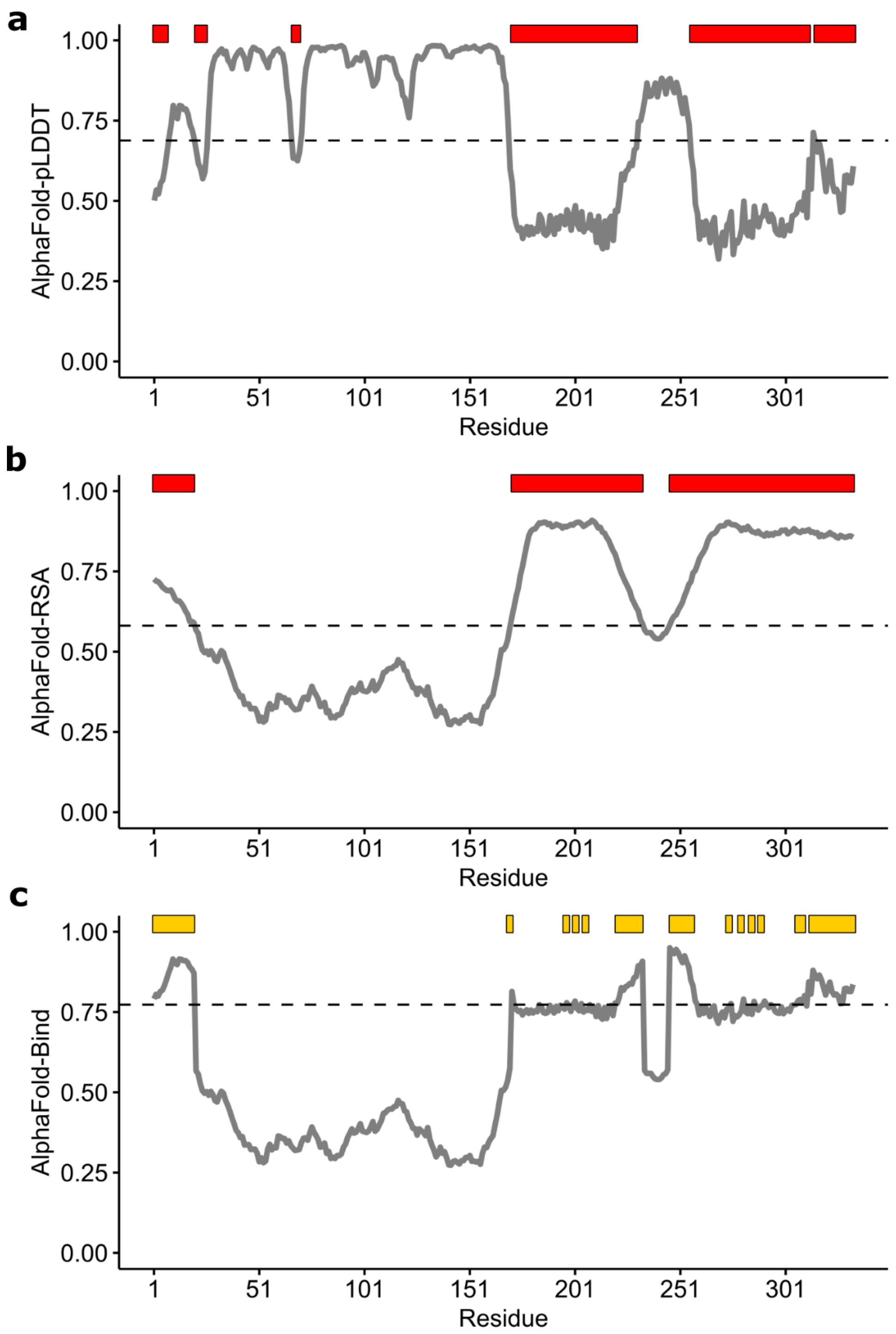


**Supplementary Figure 1 - Intrinsic disorder and conditional folding per-residue scores derived from AlphaFold prediction.** AlphaFold-pLDDT **(a)**, AlphaFold-RSA **(b)** and AlphaFold-Bind **(c)** scores and optimal thresholds for the human Ephrin-B2 protein (UniProt accession: P52799). Horizontal bars indicate sequence positions passing the threshold (dashed line). Optimal thresholds are 0.688 (AlphaFold-pLDDT), 0.581 (AlphaFold-RSA) and 0.773 (AlphaFold-Bind).
